# Automated Threshold Selection for Cryo-EM Density Maps

**DOI:** 10.1101/657395

**Authors:** Jonas Pfab, Dong Si

## Abstract

Recent advances in cryo-EM have made it possible to create protein density maps with a near-atomic resolution. This has contributed to its wide popularity, resulting in a rapidly growing number of available cryo-EM density maps. In order to computationally process them, an electron density threshold level is required which defines a lower bound for density values. In the context of this paper the threshold level is required in a pre-processing step of the backbone structure prediction project which predicts the location of Cα atoms of the backbone of a protein based on its cryo-EM density map using deep learning techniques. A custom threshold level has to be selected for each prediction in order to reduce noise that could irritate the deep learning model. Automatizing this threshold selection process makes it easier to run predictions as well as it removes the dependency of the prediction accuracy to the ability of someone to choose the right threshold value. This paper presents a method to automatize the threshold selection for the previously mentioned project as well as for other problems which require a density threshold level. The method uses the surface area to volume ratio and the ratio of voxels that lie above the threshold level to non-zero voxels as metrics to derive characteristics about suitable threshold levels based on a training dataset. The threshold level selection was tested by integrating it in the backbone prediction project and evaluating the accuracy of predictions using automatically as well as manually selected thresholds. We found that there was no loss in accuracy using the automatically selected threshold levels indicating that they are equally good as manually selected ones. The source code related to this paper can be found at https://github.com/DrDongSi/Auto-Thresholding.

## Introduction

In the last decade, advances in cryo-electron microscopy (cryo-EM) have made it possible to capture three-dimensional (3D) images of proteins with a near-atomic resolution [1]. In contrast to other approaches, such as X-ray crystallography, cryo-EM can capture these images without a crystallization of the protein [2]. Since this makes it much less costly, cryo-EM has gained wide popularity resulting in a continuously growing number of available cryo-EM protein density maps [3]–[6]. When a protein is scanned using cryo-EM the resulting image is a 3D electron density map usually stored in an MRC file format. It represents the volume of the protein complex through a 3D grid of voxels [7]. Each voxel stores the electron density value, meaning the probability that an electron is present, for its location. An electron density threshold level (later as “threshold”) is required to computationally process the cryo-EM density map. The threshold level defines a lower bound for processing the electron density voxels in a map. Voxels with value below that lower bound are set to zero [8]. Therefore, by specifying a threshold level, we can reduce experimental noise significantly. This allows researchers to focus on the structural features the map presents at a certain level (Figure 1).

**Figure 1:**
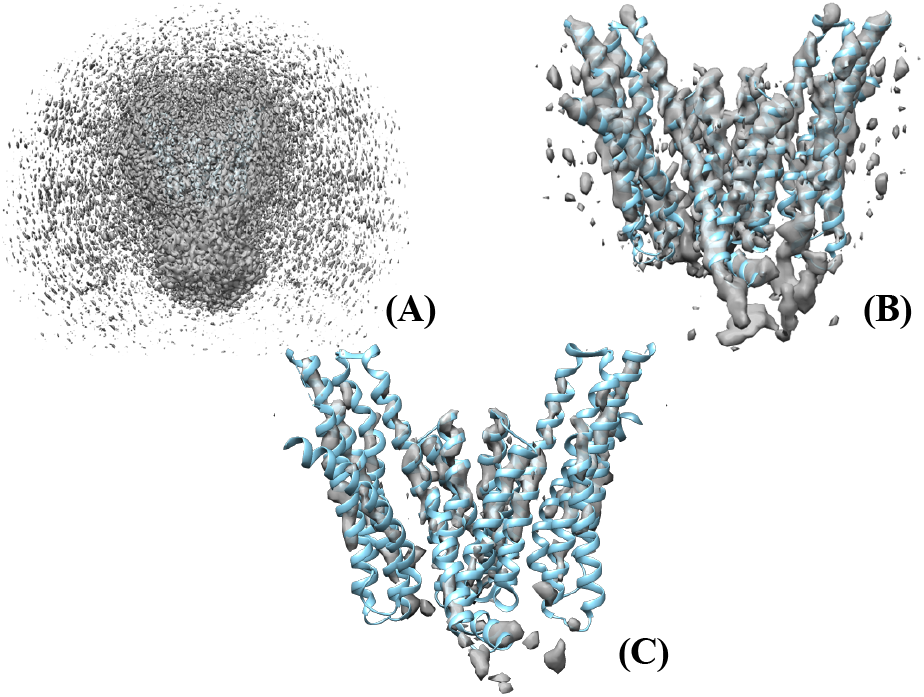
Density map EMD 8637 (gray density volume) and corresponding protein structure PDB 5v6p (cyan ribbon) for different threshold values (A) Low threshold value (B) Medium threshold value (C) High threshold value

There are several projects where the explicit selection of a threshold level is necessary [9]–[15]. In the context of this paper the threshold level is required in a pre-processing step for predicting the backbone protein structure from an electron density map [16]. In contrast to other methods, such as Rosetta and Phenix [17], [18], this project utilizes a deep learning model for the prediction [16]. All voxels below the threshold level are set to zero in order to remove noise that could irritate the deep learning model. Unfortunately, we cannot use the same threshold for different density maps since the range and distribution of electron density values varies significantly from map to map. Therefore, we have to specify a custom threshold level for each prediction [16]. And choosing the right level is crucial since it has a great impact on the accuracy of the final prediction. In Table 1 we can see the prediction results for the same density map and different threshold levels as shown in Figure 1.

**Table 1:** Backbone Prediction Results for EMD 8367 density map with varying thresholds

| Threshold Level | % of Ca Predicted | RMSD | # Incorrect |
| --- | --- | --- | --- |
| 0.005 | 86.85 | 1.69 | 50 |
| 0.035 | 76.29 | 1.58 | 10 |
| 0.130 | 00.00 | ∞ | 0 |

The obvious benefit of automating the threshold level selection is that it requires less work to run predictions. In particular, for batch predictions it can be tedious work to look at each density map individually to determine a suitable threshold level. However, there can also be a qualitative advantage using an automated selection process. If selected manually, we would choose threshold levels such that the density maps look similar to previous predictions with good results. This is problematic as it is a highly subjective process that depends on the experience of the person choosing the threshold level and therefore, is prone to errors. Particularly, if someone does not have any experience through previous predictions the selection process is essentially random. One could of course try multiple threshold levels and see which one would achieve the best results. However, since there is a large number of possible levels and each prediction is computationally very expensive this is not a viable option in most cases. Therefore, it is important that the threshold level selection process is automated. In Chimera [8] this is done by automatically selecting a threshold level, such that 1% of all voxels lie above it [19]. This method, however, is not suitable for the backbone prediction project as the total number of voxels can vary for the same density map since the size of the bounding box can change, resulting in different threshold levels for the same density map. Therefore, this paper proposes a new method to automate the threshold level selection process.

The theory of this method is presented in the Methods section. In the Results section we assess the accuracy of the automatically selected thresholds by evaluating the accuracy of backbone predictions that use automatically selected thresholds. The Discussion section examines the implications of the results and what they mean for potential other projects. Finally, in the Conclusion, we recapture our findings.

## Methods

In this section we present a method to automatically select the threshold level for a given density map. First, we talk about the general idea behind the selection process, and then we continue to show the specific methods we use. It is important to note that everything presented in this section can be used for any purpose where a threshold level has to be selected. Only in the Results section, we use the method specifically for the backbone prediction project.

The goal of this paper is to find a function *f*: *D* → ℝ, which maps any density map *d* ∈ *D* to a threshold value *t* ∈ ℝ which is suitable for *d*. We say that a threshold level is suitable for a density map if it fulfills the properties desired for a problem. In the context of the backbone prediction project this would mean that a threshold level is suitable for a density map if it results in an accurate prediction of the backbone structure.

In order to find the function *f*, we require a training set *T* consisting of tuples (*d*, *t*_*d*_) where *d* is a density map and *t*_*d*_ is a manually selected suitable threshold level for *d*. Next, we need to have some metric with which we can measure any density map *d* for a given threshold value *t*. Let *m*(*d*, *t*) be the function that implements this measurement. We can evaluate *m*(*d*, *t*_*d*_) for each density map *d* and their manually selected threshold *t*_*d*_ to find out which measured value indicates that the threshold is suitable. We can calculate an overall target metric value *m*_*t*_ using equation 1 for which, if measured for a certain threshold, we assume that it is suitable. Once we know *m*_*t*_ we can calculate a threshold level for a new density map *d* by finding the root *t* of the function *g* shown in equation 2. This is accomplished using the **root scalar** method from the SciPy library [20].

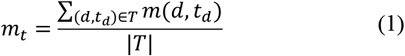

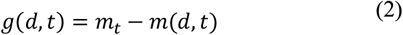

If we want to use a target metric value to find a suitable threshold level the function *m*(*d*, *t*) has to fulfill the following three properties.

1. The result of the function has to be dependent on the threshold value *t*
2. The function has to be a monotonic function so that *g* has only one root
3. For different density maps *d* and their suitable thresholds *t*_*d*_ the deviation of *m*(*d*, *t*_*d*_) should be minimal

Now, that we have developed the basic model for the threshold level selection we can look at metrics that we can use to build it. The first metric we look at is the surface area to volume ratio (SA:V). It describes the relationship between the surface area of the density map to the volume encapsulated by the surface. In general, the SA:V ratio decreases as an object gets larger and increases when it gets smaller or thinner. We can measure both attributes using the chimera commands **measure volume** and **measure area** [21]. We define *m*_*sav*_(*d*, *t*) as the function measuring the SA:V ratio. In order to validate that this function fulfills the requirements listed above we plot it on a graph for a set of density maps for which we know an ideal threshold level. The plots from a sample of five density maps are shown in Figure 2. As we can see, the SA:V ratio is dependent on the threshold level t. It also increases monotonously for growing threshold levels, as the density map becomes smaller. Finally, we can note that the SA:V ratios are relatively similar around the manually selected threshold although there are some variations. Therefore, *m*_*sav*_(*d*, *t*) is a valid candidate for the threshold level selection.

**Figure 2:**
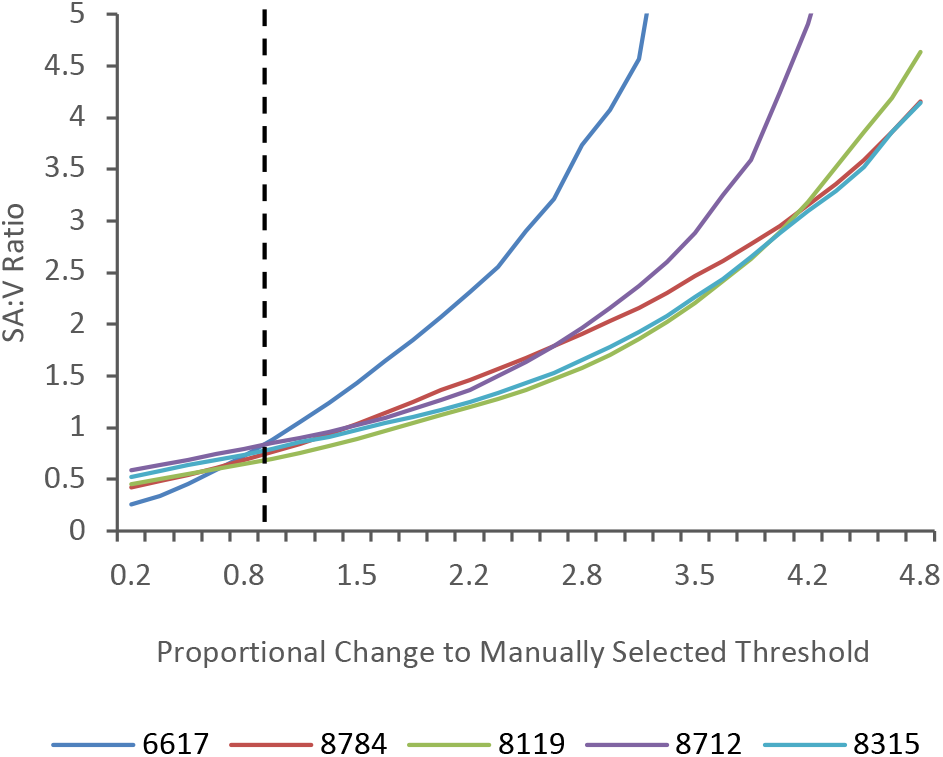
Plot of SA:V ratio for set of density maps for which we manually selected a threshold level. The x-axis shows the proportional change of the threshold level used to calculate the SA:V ratio compared to the manually selected threshold level. The vertical black line depicts the manually selected threshold level.

As we saw in Figure 2, the SA:V ratio varies only slightly around the manually selected threshold level for all density maps. However, the variations are not neglectable and might cause the threshold level prediction to be inaccurate. Therefore, we introduce another metric, the R:NZ ratio, to predict a second threshold level, with the goal of minimizing such inaccuracies. The final threshold level prediction is then derived from a weighted average of both threshold levels. The weights are calculated by solving the equation *Aw* = *b*, where *A* is a matrix in which each row contains the thresholds that were predicted for a density map from the training set using both metrices, and *b* is a vector containing the manually selected thresholds. The vector *w* that solves the equation contains the weights for each metric and is approximated using the **least-squares** function from the SciPy library [22]. The R:NZ (remaining to non-zero) ratio describes the ratio of electron density values larger than the threshold level, to electron density values larger than zero. Similarly to the SA:V ratio, we define *m*_*rnz*_(*d*, *t*) as the function measuring the R:NZ ratio and plot it on a graph (Figure 3) for a set of density maps for which we manually selected a threshold level. We can again note that the ratio is dependent on the threshold level *t* and that it decreases monotonically. Around the manually selected threshold level there are again some variations, however, they are within a small range.

**Figure 3:**
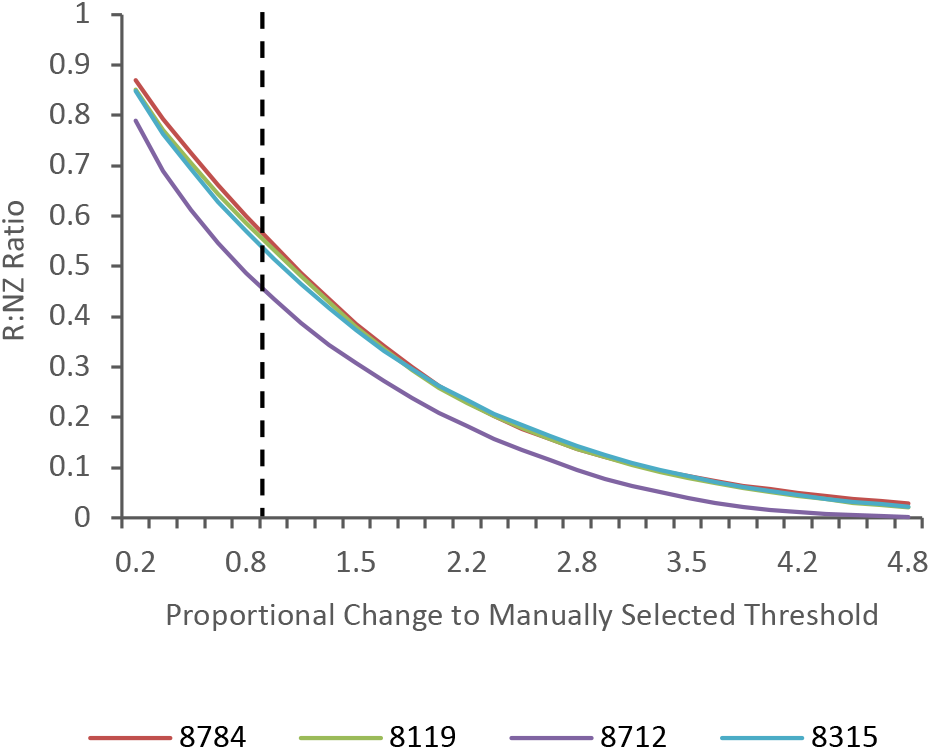
Plot of R:NZ ratio for set of density maps for which we manually selected a threshold level. The x-axis shows the proportional change of the threshold level used to calculate the R:NZ ratio compared to the manually selected threshold level. The vertical black line depicts the manually selected threshold level.

Now, that we know which metrics to use we can apply the automatic threshold level selection as following. First, we need to have a set of density maps for which we manually selected a suitable threshold level. Next, we use equation 1 to calculate the target metric value for both, the SA:V and R:NZ ratio. In order to predict the threshold level for a new density map we find the threshold levels for which both ratios equal their target metric value and calculate the average of those two levels. The result of this is the final threshold level prediction.

## Results

In order to test the accuracy of the automatic threshold prediction method presented in the previous section, we integrate it into the prediction pipeline of the deep learning based protein backbone structure prediction project. Its goal is to predict the Cα atoms of the backbone structure of a protein based on its cryo-EM density map. A more detailed description of the project can be found in [16]. The threshold selection is executed as part of the pre-processing and abolishes the need to specify custom threshold levels for each prediction.

The target metric values of the SA:V and R:NZ ratios are calculated by applying equation 1 on a training set of 27 different density maps for which we manually select a threshold level which results in an accurate backbone prediction (this training set is unrelated to the one used to train the deep learning model). The resulting target metric values are 0.9331 for the SA:V ratio with a weight of 0.26 and 0.4453 for the R:NZ ratio with a weight of 0.74. Now that we know these values we are able to integrate the threshold level prediction into the project and apply the Cα backbone structure prediction on density maps without manually specifying a threshold level. We measure the accuracy of a backbone prediction through three different metrics. The RMSD value, which describes the average distance between atoms of the predicted backbone structure and the true backbone structure, the percentage of Cα atoms of the true backbone structure which are within 3Å of Cα atoms of the predicted backbone structure, and the number of Cα atoms which are predicted incorrectly [16]. The results of the backbone predictions for a validation set of 29 different density maps using automatically and manually selected thresholds can be seen in Table 2.

**Table 2:** Protein backbone structure prediction results using both manually and automatically selected threshold levels. % Cα in 3Å describes the percentage of Cα atoms from the ground truth structure that are within 3Å of a predicted Cα atom. The RMSD value expresses the average distance between atoms of the ground truth and predicted backbone structure. The # Incorrect column shows the numbers of Cα atoms that were predicted incorrectly. The error column shows the relative error of the automatically selected threshold compared to the manual threshold and is calculated by dividing the absolute difference of both thresholds by the manual threshold value.

| EMDB ID | Threshold Level | | | % $\text{Ca}$ in 3Å | | RMSD | | # Incorrect | |
| --- | --- | --- | --- | --- | --- | --- | --- | --- | --- |
|  | Automatic | Manual | Error | Automatic | Manual | Automatic | Manual | Automatic | Manual |
| 3121 | 0.031 | 0.030 | 0.033 | 81.95 | 82.65 | 1.439 | 1.436 | 228 | 242 |
| 5623 | 0.124 | 0.110 | 0.127 | 96.04 | 96.92 | 1.034 | 1.036 | 55 | 40 |
| 5995 | 0.013 | 0.017 | 0.235 | 96.60 | 93.39 | 1.075 | 1.090 | 53 | 65 |
| 6272 | 0.010 | 0.009 | 0.111 | 99.75 | 99.74 | 0.846 | 0.853 | 0 | 0 |
| 6324 | 0.011 | 0.015 | 0.267 | 84.95 | 77.72 | 1.242 | 1.207 | 77 | 85 |
| 6346 | 5.891 | 6.000 | 0.018 | 94.91 | 94.61 | 1.239 | 1.213 | 15 | 29 |
| 6408 | 1.409 | 0.550 | 1.562 | 90.35 | 96.47 | 0.953 | 0.957 | 27 | 18 |
| 6488 | 0.023 | 0.025 | 0.080 | 88.68 | 88.79 | 1.154 | 1.135 | 276 | 288 |
| 6551 | 0.027 | 0.020 | 0.350 | 96.39 | 95.87 | 1.199 | 1.241 | 19 | 33 |
| 6617 | 0.055 | 0.070 | 0.214 | 82.79 | 78.43 | 1.098 | 1.112 | 1053 | 948 |
| 6676 | 0.017 | 0.025 | 0.320 | 80.04 | 73.44 | 1.092 | 1.050 | 78 | 51 |
| 6770 | 0.044 | 0.032 | 0.375 | 86.89 | 90.54 | 1.270 | 1.329 | 66 | 83 |
| 7063 | 0.034 | 0.025 | 0.360 | 94.86 | 96.34 | 1.057 | 1.104 | 61 | 124 |
| 8015 | 0.036 | 0.040 | 0.100 | 98.92 | 98.84 | 0.916 | 0.911 | 11 | 4 |
| 8118 | 3.240 | 3.500 | 0.074 | 93.53 | 93.21 | 1.080 | 1.049 | 36 | 37 |
| 8119 | 3.408 | 2.500 | 0.363 | 87.88 | 90.67 | 1.255 | 1.224 | 32 | 20 |
| 8289 | 0.025 | 0.020 | 0.250 | 74.87 | 80.80 | 1.557 | 1.618 | 93 | 153 |
| 8315 | 0.015 | 0.012 | 0.250 | 80.11 | 84.08 | 1.381 | 1.413 | 44 | 124 |
| 8331 | 0.033 | 0.040 | 0.175 | 96.04 | 94.30 | 1.104 | 1.086 | 213 | 165 |
| 8354 | 0.029 | 0.030 | 0.033 | 96.49 | 95.62 | 1.047 | 1.054 | 50 | 46 |
| 8482 | 0.400 | 0.270 | 0.481 | 67.55 | 79.79 | 1.577 | 1.586 | 27 | 31 |
| 8515 | 2.847 | 2.000 | 0.424 | 94.68 | 96.67 | 1.091 | 1.146 | 4 | 10 |
| 8642 | 0.037 | 0.020 | 0.850 | 86.95 | 90.21 | 1.370 | 1.383 | 22 | 171 |
| 8651 | 1.829 | 1.400 | 0.306 | 75.09 | 82.25 | 1.585 | 1.527 | 10 | 15 |
| 8712 | 0.010 | 0.010 | 0.000 | 89.67 | 89.87 | 1.192 | 1.160 | 6 | 9 |
| 8764 | 0.014 | 0.023 | 0.391 | 92.42 | 78.19 | 1.135 | 1.096 | 107 | 71 |
| 8782 | 1.128 | 1.000 | 0.128 | 84.57 | 88.15 | 1.433 | 1.445 | 50 | 45 |
| 8881 | 0.022 | 0.010 | 1.200 | 91.08 | 84.96 | 1.226 | 1.216 | 64 | 17 |
| Avg. |  |  | **0.324** | **88.72** | **89.02** | **1.202** | **1.203** | **96** | **104** |

In order to examine the differences between manually and automatically selected thresholds more closely, we take a look at an example, the prediction of the EMD 6551 density map (Figure 4). We can see the prediction results (tan color) using the manually selected threshold in (A) and the prediction results using the automatically selected one in (B). Both predictions are compared to the true backbone structure shown in pink color and embedded in the density map at the threshold level that was used for each prediction. The automatically selected threshold level (0.027) was higher than the manually selected one (0.02) which, on average, results in a lower number of predicted Cα atoms, as the number of voxels with non-zero density values decreases. Since the thresholds differ only slightly this is barely noticeable when looking at the complete backbone structures. However, if we zoom into some areas we can detect minor differences. In the enlarged area on the upper right of both predictions we can see that, for the manually selected threshold, some Cα atoms were predicted incorrectly in locations where there are no Cα atoms in the true structure. In contrast to that we can see that in the enlarged area on the center left of both predictions, for the prediction which used the automatically selected threshold level, some Cα atoms are missing where they were correctly predicted for the manually selected threshold. Therefore, we cannot conclude that one prediction is better than the other solely through visual examination.

**Figure 4:**
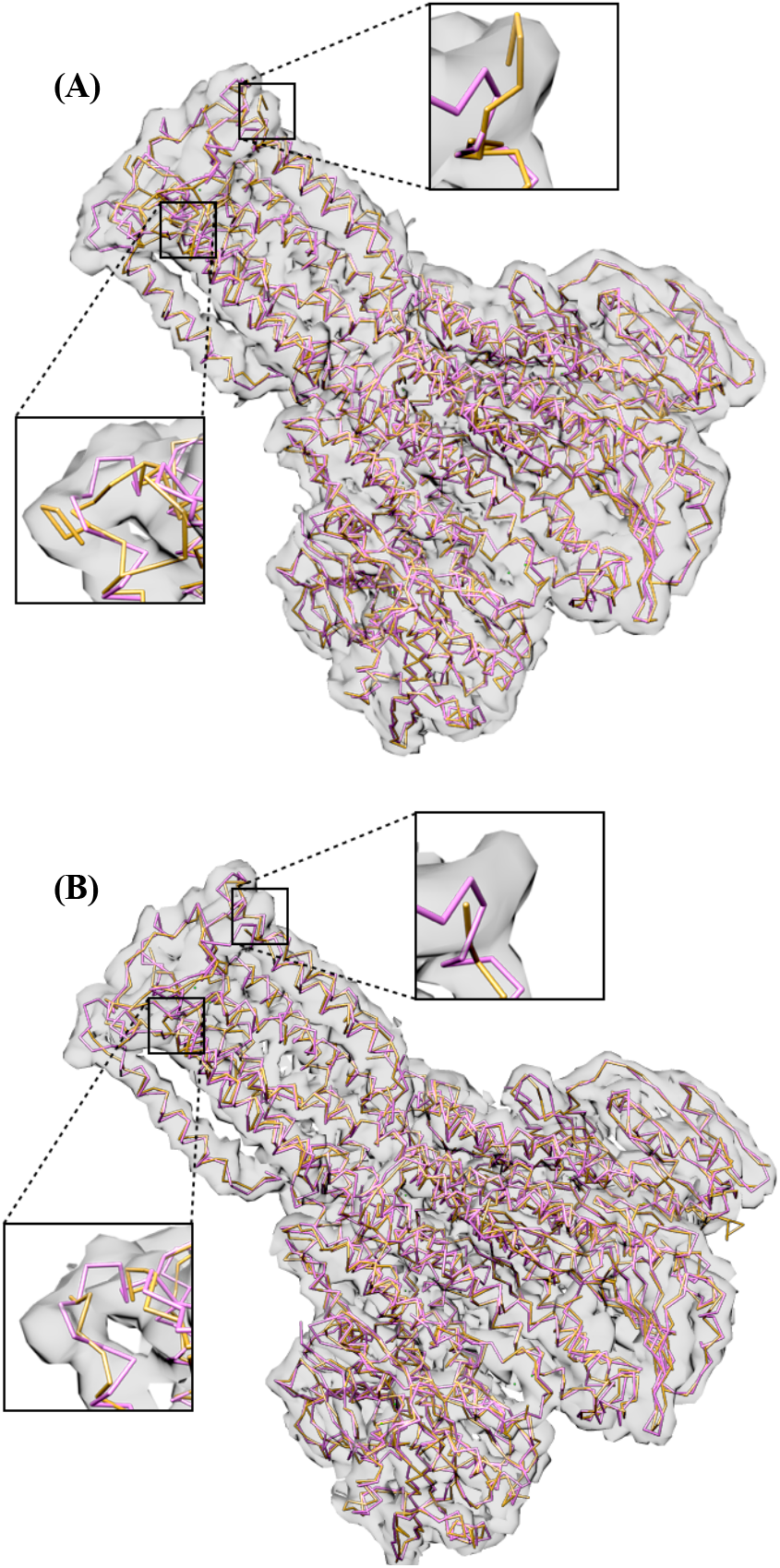
Backbone structure prediction results (tan color) for EMD 6551 density map, using automatically and manually selected threshold levels, compared to the true backbone structure (PDB 3jcf) in pink color. The structures are embedded in the density map at the threshold level used for the prediction (A) Prediction result using manually selected threshold level of 0.02 (B) Prediction result using automatically selected threshold level of 0.027

## Discussion

In this section, we evaluate the implications of the results, discuss how the automated threshold prediction can be used in other projects, and talk about possible future work.

The results from Table 2 show that the automatic threshold prediction method is able to select thresholds with a mean relative error of 0.32 compared to the manually selected thresholds, yielding backbone structure predictions of similar accuracy. Although the average percentage of Cα atoms which are within 3Å of their predicted counterpart decreased by 0.3%, this is only a minor difference which stands in contrast to the average RMSD value and number of incorrectly predicted atoms which both improved using the automatically selected threshold levels. Therefore, we can say that the automatic threshold prediction can be integrated in the backbone prediction project to abolish the need for a manual selection without a loss in prediction accuracy. This makes it easier to execute a backbone prediction, in particular, for researchers who are not familiar with the underlying logic behind the prediction.

Besides its application in the backbone prediction project, the goal of the automatic threshold selection method was that it can be applied to other problems, where a threshold level is required, as well. Unfortunately, we cannot give a universal answer about whether or not this is achieved, since it depends on the specific characteristics of the problem. We can, however, outline a general property about suitable threshold levels for a certain problem, which must be true in order for the automatic threshold selection to be useful. This is that the SA:V and R:NZ ratios have to be similar for different density maps and their suitable threshold levels. If this is not the case, we cannot find a meaningful metric target value for either ratios for which, if measured for a certain threshold, we can assume that it is suitable. The benefit of our proposed method is that it can easily be customized for a certain problem. Therefore, one can evaluate whether or not it works for that problem with little effort.

Future work might include finding new metrics which could be used to predict the threshold levels. This could then easily be added to the existing method and further minimize the inaccuracy of threshold predictions. Another approach to improve the threshold selection might be to apply machine learning techniques. This would potentially, however, include extensive pre-processing and training efforts which would make it significantly more difficult to customize the threshold selection to different problems.

## Conclusion

In summary, we presented a method to automatically select density threshold levels that are required in order to process cryo-EM density maps e.g. for visualization and modelling. We integrated the automatic selection process into the backbone structure prediction project [16] without a loss in accuracy compared to manually selected threshold levels. For a validation set of 29 density maps the average RMSD value even improved from 2.03 to 1.202 and the average number of incorrectly predicted Cα atoms decreased from 104 to 96. The automatic threshold selection further automatizes the prediction process, as well as it removes the dependency of the prediction accuracy to the ability of someone to choose the right threshold value. We developed the method such that it can easily be customized for other problems which require the selection of a threshold level, through a simple training process. To calculate the threshold level for a density map we utilized the surface area to volume ratio as well as the ratio of voxels above the threshold to non-zero voxels. These metrics were chosen since they resulted in similar values for different density maps at their suitable threshold. Further research could encompass the addition of further metrics to increase the selection accuracy, as well as the application of machine learning techniques to solve the threshold prediction problem.

## Acknowledgments

We gratefully acknowledge the support of NVIDIA Corporation (Santa Clara, CA, USA) with the donation of the GPU used for this research. This research was funded by the Graduate Research Award of Computing and Software Systems division and the startup fund 74-0525 of the University of Washington Bothell.

